# Fundamental metabolic strategies of heterotrophic bacteria

**DOI:** 10.1101/2022.08.04.502823

**Authors:** Matti Gralka, Shaul Pollak, Otto X. Cordero

## Abstract

Through their metabolism, heterotrophic microbes drive carbon cycling in many environments (1). These microbes consume (and produce) hundreds to thousands of different metabolic substrates, begging the question of what level of description is required to understand the metabolic processes structuring their communities: do we need to account for the detailed metabolic capabilities of each organism, or can these capabilities be understood in terms of a few well-conserved carbon utilization strategies that could be more easily interpreted and more robustly predicted? Based on the high-throughput phenotyping of a diverse collection of marine bacteria, we showed that the fundamental metabolic strategy of heterotrophic microbes can be understood in terms of a single axis of variation, representing their preference for either glycolytic (sugars) or gluconeogenic (amino and organic acids) carbon sources. Moreover, an organism’s position on this axis is imprinted in its genome, allowing us to successfully predict metabolic strategy across the bacterial tree of life. Our analysis also unveils a novel and general association between metabolic strategy and genomic GC content, which we hypothesize results from the difference in C:N supply associated with typical sugar and acid substrates. Thus, our work reveals a fundamental constraint on microbial evolution that structures bacterial genomes and communities and can be leveraged to understand diversity in functional terms, beyond catalogs of genes and taxa.

---

Next-generation sequencing has enabled the collection of enormous amounts of data about the abundances of taxa or genes in microbial communities (2–4). However, outside of highly specialized and taxonomically conserved functions like photosynthesis or ammonia oxidization, it often remains difficult to associate taxa to the processes that take place in a community. This difficulty is particularly acute in heterotrophic microbial communities, which perform many important ecosystem functions such as organic matter recycling, mediated by the metabolic actions and interactions of their members. This is in part because most bacterial species are not experimentally characterized and because we lack the ability to accurately predict phenotypes from genomes. A more fundamental problem, however, is that the functional space of heterotrophic metabolism is enormous (5), given the hundreds to thousands of metabolites typically found in heterotrophic ecosystems that each can serve as potential carbon and energy source for some subset of the community (6).This substrate diversity gives rise to a huge number of potential metabolic niches, about 10^30^ for just 100 metabolites. Therefore, solving the challenge of understanding metabolic processes in heterotrophic microbial communities requires not only experimental characterizations and the development of phenotype-genotype mappings, but also a way to conceptually simplify, i.e., to “coarse-grain”, metabolic niche space into a few easily interpretable metabolic strategies (7, 8).

We approach this problem through the high-throughput metabolic profiling of 186 strains of marine heterotrophic bacteria. Most strains were originally isolated from communities growing on polysaccharide particles (9, 10) and chosen to maximize diversity across five abundant marine orders, while also including close relatives to span a wide range of pairwise phylogenetic differences (SI Fig. S1, SI Table 1). All species were grown individually in minimal media containing one of 135 potential carbon substrates, ranging from glucose to amino acids to various polysaccharides (SI Table 2, see methods), for 15 days. Growth over time was assessed by measuring optical density at least once per day and quantified by extracting growth rates, lag times, and yields from logistic fits (Fig 1a).

**Fig. 1.**
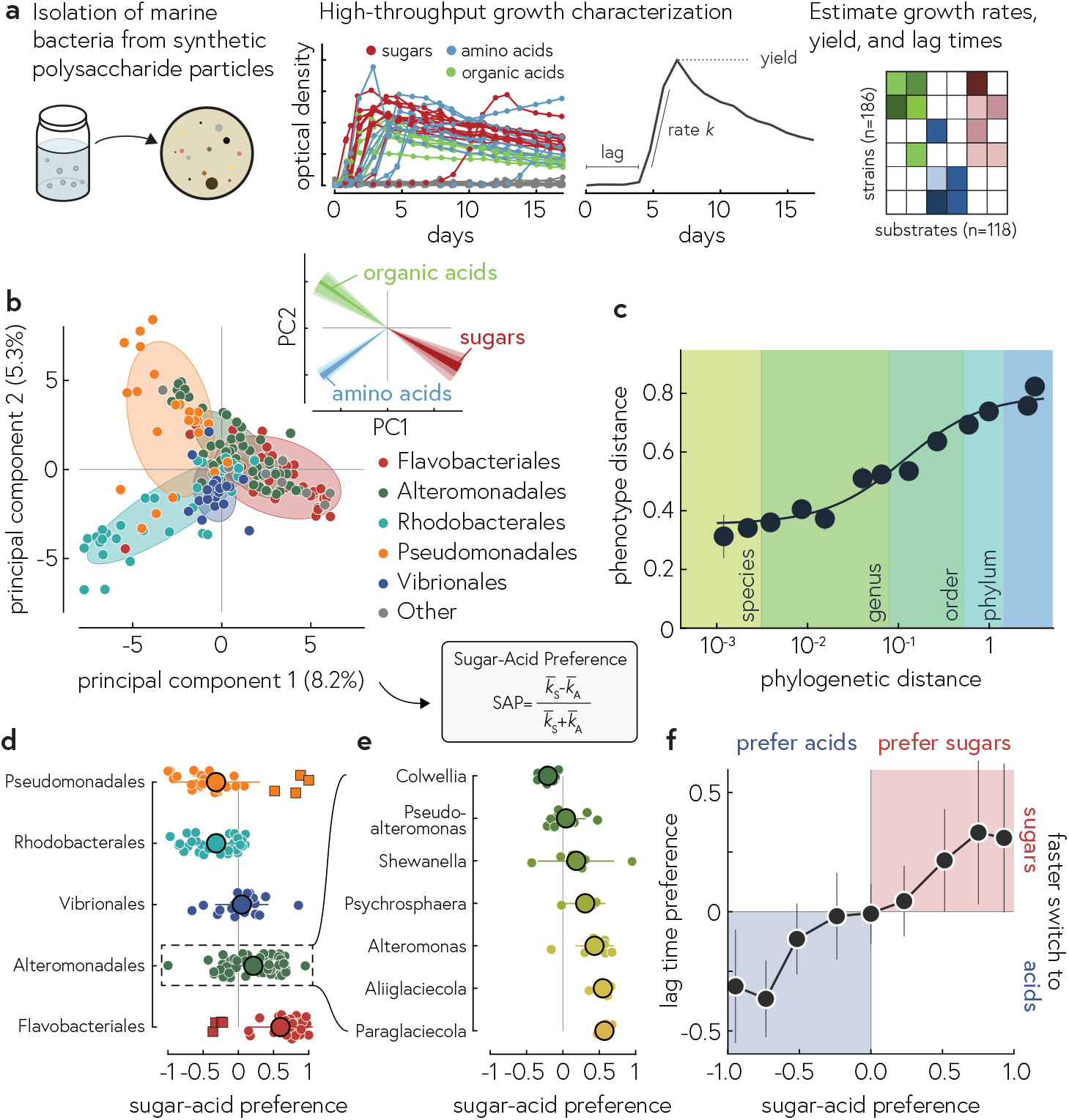
a) Outline of the experimental procedure: 186 marine isolates were grown in high throughput over 17 days on 118 individual substrates, including various sugars, organic acids, and amino acids. We characterized the resulting growth curves in terms of the lag time, growth rate, and yield. b) Principal component analysis of the growth rates revealed overall taxonomic cohesion and similarity between different taxonomic groups. Averaging the loadings (how much each substrate contributes to the first and second principal component, inset; see SI Fig S5 for the individual loadings) provided a biological interpretation for the first principal component as the preference for sugars or acids (inset), which we quantify as the sugar-acid preference (SAP). c) The phenotypic distance (cosine distance between growth vectors) was correlated with the phylogenetic distance (from GTDB-tk, see SI Fig. S8 for other measures of genomic similarity) but reached neither 0 nor 1 for very closely or distantly related strains, respectively (vertical lines correspond to the average phylogenetic distance at the different taxonomic levels). Thus, there was in general no characteristic phylogenetic distance below which we can expect all strains to have the same phenotype. d) Sugar-acid preference was variable between species but roughly conserved at the order level; exceptions to the order-wide tendencies (squares) can be explained by fundamental niches that differ from the order median (see main text). e) Variability within orders was mostly explained by preferences at the genus level, but some genera were highly variable within themselves (e.g., *Shewanella*). Only genera within Alteromonadales with at least three representatives are shown. f) Correlating sugar-acid preference with the preference in switching rate to sugars or acids revealed that different characteristics of the growth are correlated. Species that prefer sugars had, on average, shorter lag times on sugars than on acids, and vice versa. Error bars represent standard deviations.

The resulting matrix of more than 22,000 growth rates forms the basis for our subsequent analysis (SI Table 3). We observed growth in 37% of all conditions, but the probability of observing growth varied widely both across strains (SI Fig. S2) and substrates (SI Fig. S3): species consumed between 2 and 66 different substrates, and substrates were consumed by between 1 (tartrate) and 155 (glucose) strains. Of the 135 substrates tested, 118 supported robust growth of at least one species. We found practically insignificant correlations between the number of consumed carbon sources, growth rate, and yield, and no evidence for a trade-off between growth rate and yield (SI Fig. S4).

To uncover patterns in the matrix of growth rates, we performed principal component analysis of the normalized (by species) growth rates (Fig. 1b) and discovered that the first principal component, i.e., the fundamental dimension of metabolic phenotypes, corresponded to species’ preferences for either acids or sugars. To show this, we averaged the loadings of each substrate according to their fundamental chemical class (sugar, amino acid, organic acid); individual loadings are shown in SI Fig. S5. To quantify this correspondence, we introduced the Sugar-Acid Preference (SAP), defined from the average growth rate on sugars *k*_*S*_ and acids *k*_*A*_ as by

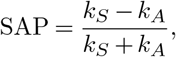

which ranged from +1 for extreme sugar specialists (no growth on any acid) to − 1 for extreme acid specialists (SI Fig. S5). Given its high correlation with the first principal component (*R*^2^ = 0.92, SI Fig. S6), we focused on the SAP as the primary index to quantify the metabolic preference of each strain; while the second principal component correlated mostly with the preference for amino or organic acids, that correspondence was weaker (*R*^2^ = 0.5, see SI Fig. S6b). Growth phenotypes and thus SAP were largely reproducible across replicate experiments, such that we used the average across up to three biological replicate experiments in the following (SI Fig. S7 and Supplementary Note 1).

More closely related strains tended to have more similar metabolic capabilities, but there was no fixed taxonomic scale or measure of phylogenetic distance (e.g., via marker genes, total gene content or pathway content, see SI Fig. S8) where metabolic preferences became coupled or uncoupled (Fig. 1c). As a result, the SAP was imperfectly correlated with taxonomy (Fig 1d): for instance, species in the order Flavobacteriales are known to degrade polysaccharides (11) and indeed, they tended to prefer sugars in our screen (SAP > 0). However, exceptions to the median metabolic preference emerged, indicated by squares in Fig. 1d, such as the acid-specialist *Tenacibaculum* genus in the Flavobacteriales, which includes fish pathogens (12). Conversely, the orders Pseudomonadales and Rhodobacterales (commonly thought to specialize in simple substrates (13)) tended to prefer acids (SAP < 0), but we also found the sugar-specialist Pseudomonadales genus *Saccharophagus*, which are known sugar degraders (14).

Beyond quantifying the preferences for growth on sugars and acids, the SAP also predicted lag time differences when switching between preferred and non-preferred substrates (Fig 1a). Recent theory and experimental results in *E. coli* and *P. aeruginosa* have suggested that different regulatory programs may prime organisms to grow rapidly on glycolytic or gluconeogenic substrates upon switching between substrate types. We defined the lag-time preference analogously to the sugar-acid preference, via the average (inverse) lag times on sugars *τ*_*S*_ and acids *τ*_*A*_, as 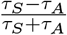. We found that sugar specialists on average tended to switch faster to growth on sugars than on acids, and vice-versa (Fig 1d). This suggests that the theory may indicate a general connection between metabolic strategies and lag times rooted in biochemical trade-offs.

## Genotype-phenotype mapping

What are the genomic underpinnings of the measured phenotypes? To solve this structure-function question and understand what aspects of the phenotypes are predictable from the genome, we connected our measured phenotype data with the annotated draft genomes. First, we identified complete substrate degradation pathways (see Methods) to predict whether a given species would be able to grow on the respective substrate. Surprisingly, we found the predictability on an individual species-substrate level to be at chance level (SI Fig S9); by any measure, growth of an individual species on a given carbon substrate cannot be predicted accurately using current state-of-the-art annotation and metabolic model creation pipelines.

While the ability to consume a specific substrate cannot be predicted from genomic information (at least not with current state-of-the-art tools), we did observe a correlation between the overall metabolic strategy (i.e., SAP) and the number of Carbohydrate-Active Enzymes (SI Fig. S10). Importantly, we observed this trend even within orders (except Vibrionales), indicating a biological signal beyond pure taxonomy. This observation also explains the “phenotypic outliers” in Fig 1d: flavobacteria with an acid preference (genus *Tenacibaculum*) have fewer than average CAZymes, whereas pseudomonads preferring sugars (genus *Saccharophagus*) have more CAZymes than expected for that clade; these genera thus occupy fundamentally different niches than the typical representative from that order in our strains. These findings led us to hypothesize that overall metabolic preferences may be predictable, even if specific growth phenotypes are not.

Consistent with the increased number of CAZymes, we found that sugar specialists tended to have more genes in sugar degradation pathways, and conversely for acid specialists. As seen in the examples of the galactose (sugar) and propionate (acid) degradation pathways (Fig 2a) and more generally in Fig. 2b, we found that the relative abundance of genes in sugar and acid catabolism pathways was, on average, positively and negatively correlated with SAP (Fig 2b), respectively.

**Fig. 2.**
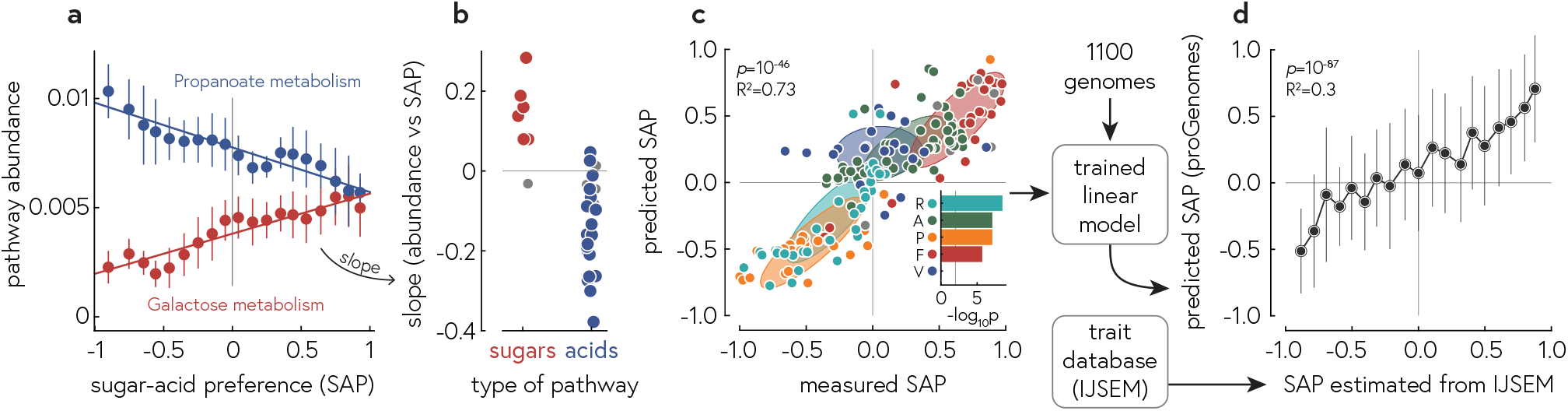
a) Two examples of pathway abundances (number of genes in the pathway normalized by the total number of genes in each strain) as a function of sugar-acid preference. b) Computing the regression slopes in panel a across all pathways, we found that sugar specialist strains tended to have more genes in sugar catabolic pathways, whereas acid specialist strains tended to have more genes in acid catabolic pathways. Gray symbols correspond to regression with a correct *p* > 0.05. c) Using the total abundance of genes in sugar and acid degradation pathways, we developed a linear model that predicted sugar-acid preference with high accuracy (*R*^2^ = 0.73), even within orders (except Vibrionales, see inset, which shows − log_10_ *p* for the linear regressions within each order). d) We estimated metabolic preferences for strains outside of our experimental study from published trait databases and contrasted these estimates with SAP prediction for species-matched reference genomes, revealing a highly significant correlation between them. Error bars represent standard deviations in panel a and standard error of the mean in c and e.

Two processes contribute to the evolution of these pathways alongside the metabolic preferences. Firstly, gaps in the central part of the pathway get filled and auxiliary functions are added, i.e., the number of unique reactions in a given pathway increases (pathway completeness). For instance, in the galactose degradation pathway (SI Fig. S12), in addition to a complete chain of enzymes converting *β*-galactose into glucose-6-phosphate, genomes with a high degree of pathway completeness have genes converting other substrates, such as galactitol or lactose, into *β*-galactose. Secondly, evolution may select for an increase in the copy number of orthologous enzymes performing a given reaction in the pathway (SI Fig. S11). These “copies” could either be bona fide duplications that simply increase gene dosage, or different variants that are likely to operate optimally under different environmental conditions. By comparing orthologs between and within genomes, we found support for the latter hypothesis, as orthologs were often more similar to genes in distantly related taxa (e.g. different phyla) than to other “copies” in the same genome. For example, the KEGG ortholog K01785, converting *β*-galactose to *α*-galactose as part of the galactose degradation pathway (SI Fig. S12), was present in up to 6 “copies” in a strain closely related to *Zobellia galactanivorans*. However, some of the gene “copies” clustered closely with genes derived from *α*- and *γ*-proteobacteria. This pattern was general, with the similarities of ortholog copies within the same organism and between organisms being statistically indistinguishable.

The correlations of pathways abundances with metabolic preferences allow us to predict phenotypes with high accuracy. We developed a generalized, 2-variable, linear model (modified to yield predictions between −1 and 1, see Methods) to predict SAP based on the total relative abundance of genes in sugar and acid degradation pathways, respectively. This model achieves remarkably high predictive power (average *R*^2^=0.73 (95% CI [0.66, 0.79]) for 1000 out-of-sample predictions, Fig 2c) and importantly, correctly predicts SAP even within taxonomic orders (Fig 2c inset) – thus, the model goes beyond simple correlations of phenotypes with taxonomy. Supplementary to this simple, yet surprisingly powerful model, a more sophisticated model based on the relative abundances of all 46 KEGG pathways in central metabolism achieves even higher predictive power (*R*^2^=0.82) and can be used where this additional power is required.

Our simple model can be used predict metabolic strategies in a wide range of bacteria, including those outside of our experimental screen. To show this, we estimated metabolic strategies from a database of bacterial traits (15). We counted the number of sugars *S* and acids *A* that each species was reported to grow on, and estimate the SAP for each species as *SAP* ≈ (*S* − *A*)*/*(*S* + *A*). Then, we matched species names with genomes in the proGenomes database (16), annotated the corresponding reference genomes, and predicted their SAP using our simple model. The result of this procedure was a highly significant (*ρ* = 0.55, *p* = 10^−87^) correlation between predicted and estimated SAP for about 1100 species (Fig. 2d), including species from two phyla (Firmicutes and Actinobacteria) for which we did not screen a single representative (SI Fig. S13). Thus, by simply quantifying the relative abundances of sugar and acid catabolic pathways, we are able to predict metabolic preferences across a wide range of bacteria.

### Connecting metabolic strategies with GC content

GC content is associated with differences in amino acid usage (17), which in turn translate to differences in the relative demand of carbon (C) and nitrogen (N): low GC organisms code for more C rich amino acids, like tyrosine and phenylalanine, while high GC organisms code for more amino acids with relatively higher N content, like glycine and arginine. As a consequence, adaptation under carbon or nitrogen limitation can bias GC content evolution (18). Accordingly, C:N supply along the oceans depth profile has been shown to correlate strongly with the average GC content of the microorganisms that live in it, such that high N supply favors high GC organisms and vice versa (19).

The idea that C:N supply could determine GC content led us to ask whether GC content could be related to the preference for sugars or acids. Sugars are rich in carbon but contain nitrogen only in rare cases (such as amino sugars in chitin). Conversely, many organic acids, especially amino acids, contain nitrogen, and those that do not (e.g., TCA cycle intermediates) have a lower carbon density than sugars. Therefore, we hypothesize that the preference for sugars or acids as defined here, should be correlated with the GC content, such that sugar specialists have a low GC and acid specialists high GC.

In agreement with this prediction, we found a highly significant correlation (*ρ* = − 0.48, *p* = 8.7 × 10^−12^) between sugar-acid preferences and genomic GC content in our experiments, such that species with low SAP (i.e., acid specialists) have an average GC of 60% whereas high SAP species (sugar specialists) have an average GC around 40% (Fig. 3b). Notably, this correlation was much stronger than the correlation with other basic characteristics of the genomes, such as the number of coding regions (*p* = 0.2). In contrast to our previous analyses of pathway abundances as a function of SAP, there was no correlation between SAP and GC within orders (SI Fig. S14), possibly because GC content evolves very slowly and is thus relatively conserved below the order level.

**Fig. 3.**
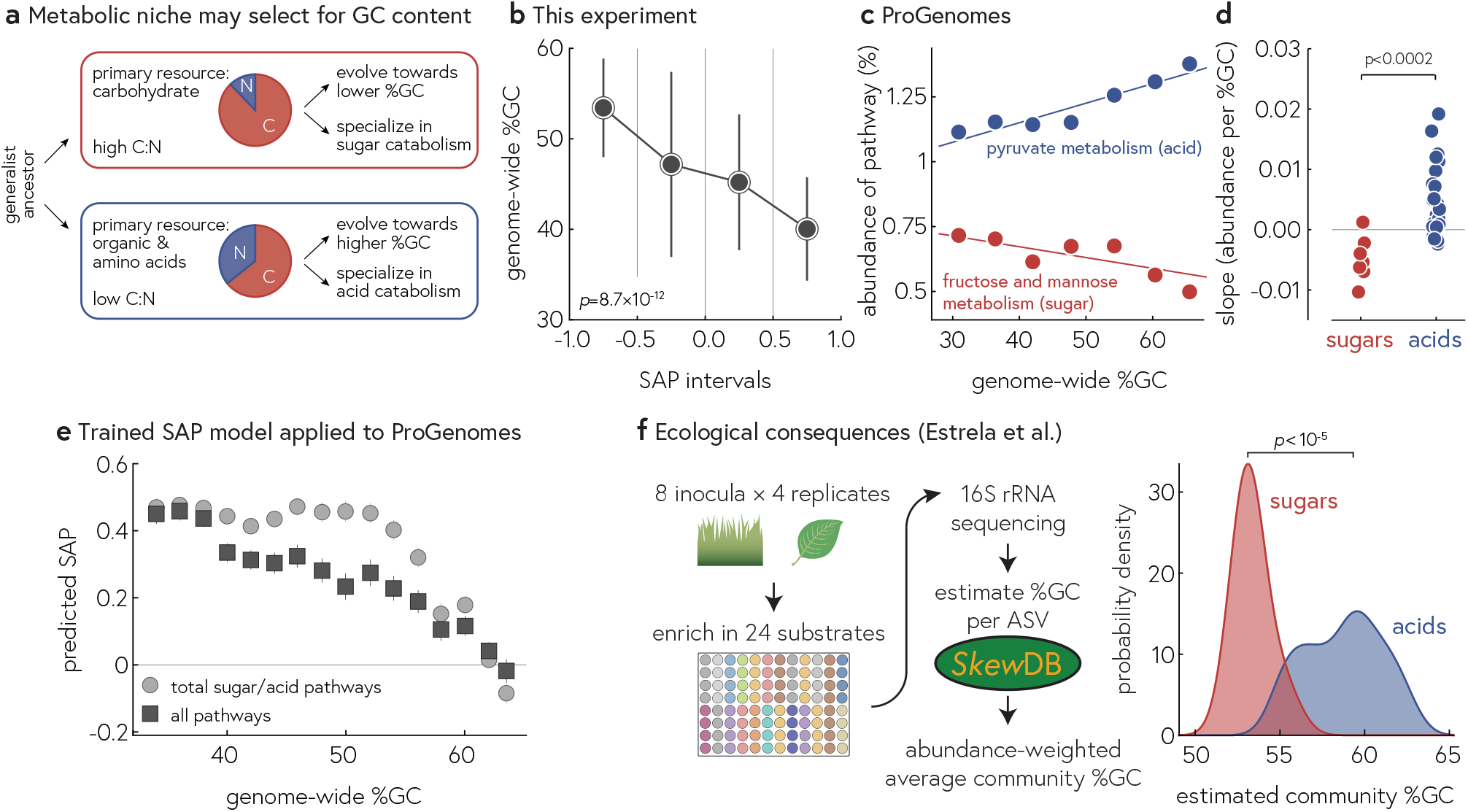
a) Description of a possible mechanism for selection for GC content in different environments: since GC content biases the nutrient requirements of the proteome, evolution carbon or nitrogen limitation can drive GC content evolution. b) In our experiment, GC content was highly correlated with sugar-acid preference. c) Example for the correlation of pathway abundance with GC content in more than 11,000 diverse reference genomes, including many phyla outside of our experimental screen. The relative abundance of genes in pyruvate metabolism increases with GC content, whereas the abundance of the fructose and mannose metabolism pathway decreases with GC. Measuring the regression slopes for all sugar/acid metabolic pathways, sugar/acid pathways tended to become less/more abundant with genomic GC. e) Predicting the SAP using our trained models (using either total abundance of sugar/acid genes or the individual abundances across all sugar/acid metabolic pathways) reveals a highly significant correlation between GC content and predicted SAP. f) Analyzing communities enriched on sugars and acids, we estimated the average community GC content, which tended to be lower for sugars than acids, in agreement with the prediction that sugar/acid specialists should have low/high GC content.

To assess whether the correlation between GC content and metabolic preferences holds more generally, we analyzed reference genomes from the proGenomes collection (11,826 unique species, 80% of which came from four large phyla (Proteobacteria, Firmicutes, Actinobacteria, and Bacteroidetes) and computed the relative abundance of central metabolic pathways. As an example, consider the fructose and mannose metabolism and the pyruvate metabolism pathways, which decreased/increased in relative abundance as a function of GC content, respectively (Fig. 3c). Extracting the linear slopes for each pathway, all of which were highly significant, we arrived at a similar picture as Fig. 2b, where sugar pathways tended to decrease and acid pathways tended to increase in abundance as a function of GC content (Fig. 3d). Consistently with this result, we predicted SAP for all reference genomes in proGenomes and found a highly significant correlation between predicted metabolic preference and GC content, i.e., acid specialists generally have higher GC content than sugar specialists (Fig. 3e). To our knowledge, this is the first description of this very general trend across a large number of genomes, sampled across the tree of bacterial life.

Finally, if the connection between SAP and GC content holds in the environment, we would expect natural microbial communities assembled on sugar substrates to display a lower community-average GC content than communities enriched on acids, and vice versa. To test this, we analyzed 16S we analyzed data from an enrichment experiment based on 8 soil inocula grown with serial passaging across 24 different carbon sources (20) (Fig. 3f). We used the database SkewDB (21) to estimate the genomic GC for each taxon in the communities and computed the abundance-weighted average community GC content. In agreement with our prediction, sugar-enriched communities had a significantly lower average GC content than acid-enriched communities (Fig 3e, TTest *p* < 10^−5^).

## Discussion

By characterizing the growth of diverse marine bacteria across many substrates, we have uncovered the preference for sugars and acids as the primary axis of metabolic strategies among marine heterotrophic bacteria (as well as heterotrophs from soil, SI Fig. S15) and identified genomic correlates that allow us to quantitatively predict these preferences.

We propose that this dichotomy of sugars and acids is rooted in the structure of central metabolism, i.e., the major pathways of glycolysis/gluconeogenesis and the citric acid cycle, and various short pathways that shuttle amino acids, organic acids, and simple sugars into these two main pathways. Recent research has indicated that biochemical constraints (e.g., the accumulation of counteracting enzymes and the resulting futile cycling) in central metabolism precludes the existence of species that grow fast on both glycolytic (e.g., glucose) and gluconeogenic (e.g., succinate) substrates, while at the same time being able to switch rapidly between both types of substrates (22, 23). Thus, we propose that species may “choose” to have a preferred direction of running their central metabolism (either glycolytic or gluconeogenic), and that this preference may be hardcoded in the genome and locks in their global metabolic strategy. Alternatively, generalists with no obvious preference for acids or sugars emerge, which may be metabolically agile but may have to pay for their metabolic capabilities by requiring high maintenance energy (22).

Our systematic exploration of metabolic strategies and their genomic correlates revealed that while coarse-grained descriptions of metabolic strategies are predictable, individual phenotypes are (currently) not. We argue that this lack of predictability is at least not entirely due to technical limitations such as incomplete annotation databases, but is an expression of the evolutionary processes at work on different time scales: the ability to degrade specific substrates can vary rapidly (on the level of species or even subspecies) because it can be switched on (e.g., through horizontal transfer of a CAZyme) or off (e.g., through gene loss or changes in regulation) very easily, even on time scales of laboratory evolution (24–26). Overall metabolic strategies evolve more slowly (perhaps at the level of genera or families) because they rely on the expansion of whole pathways, which may allow an organism to run those pathways efficiently under various environmental conditions (SI Fig. S12).

On long evolutionary timescales (at or above the level of orders), slow adaptation of genomic GC content may optimize whole taxonomic clades to the nutrient availabilities in that clade’s typical ecological niche. Thus, metabolic strategies that are well-adapted to the nutrient requirements of the proteome may be one of many potential factors shaping GC content evolution, including biases in mutations and their repair (27–29), DNA stability (30), and GC-biased gene conversion (gBGC) (31).

The dataset we have presented here constitutes, to our knowledge, the largest and most detailed metabolic characterization of bacteria from the same ecosystem to date. Datasets such as this will serve as a resource for future studies of bacterial physiology and its genomic underpinnings. By developing better genotype-phenotype mappings, we will be able to assign function to taxa, better understand metabolic interactions in microbial communities, and develop more detailed ecosystem models.

## Methods

### Media

All experiments were performed in liquid culture using either rich (MB 2216, Difco) or minimal media (*MBL*, see SI Table 2) with a substrate concentration of 30mM carbon, except for polysaccharide cultures. For a full list of substrates, see SI Table 2.

### Strains

For a full list of strains including their taxonomy and isolation details, see SI Table 1. Apart from a few exception (notably, the marine model bacterium *Ruegeria pomeroyi* DSS-3 (32) and the Vibrio strains YB2 (33), 1A06, 12B01, and 13B01), all strains were isolated from enrichments of coastal seawater on various polysaccharides. Strains were originally double-streaked at the time of isolation, and double-streaked again before being arrayed in two 96 well plates and stored in 5% DMSO at −80^deg^C.

### Phenotyping experiments

Before the main phenotyping experiment, the strain library was thawed and inoculated 1:10 into rich media. After 3 days, cultures were diluted 1:10 into minimal media without carbon for 2h, and then diluted 1:15 into prefilled 384 well plates (5µl into 75µl) containing minimal media with a single carbon source, in duplicate. Plates were sealed with MicroAmp Optical Adhesive Film (Applied Biosystems) to prevent contamination.

Bacterial growth was estimated by optical density at 600nm, which was measured at least once a day (twice a day initially) on a Tecan Spark plate reader with a stacker module (Tecan Trading AG, Switzerland). Plates were stored at room temperature in the dark in between readings.

After the experiment, growth curves were extracted and, if they crossed a minimum threshold of optical density, fitted using a custom fitting algorithm, based on the logistic growth model, that automatically estimated growth rate (the maximal exponential rate), lag time (the time before maximum growth is reached), yield (the maximal optical density reached), and death rate (the rate of decay of optical density after the maximum was reached). The R^2^ of the nonlinear fitting procedure was on average 0.9931 (CI 95% [0.9767, 0.9996]).

### Replicability

Two prior experiments were performed similarly (v1, v2) except with small alteration to inoculation protocol, carbon sources, and strains. For all three experiments, we computed the SAP, which were well-correlated across experiments (see SI Fig S7); in the main text, we report the average SAP per strain across all experiments that contained that strain. See Supplementary Note 1 for details.

### Comparison with Kehe et al

Kehe et al. (34) grew 20 soil bacteria on 33 carbon substrates for 72 hours and reported the growth yield for each condition normalized by the maximal yield per strain. We extracted the yields from their Supplementary Figure S2 using automated image analysis (replotted as SI Fig. S15a) and performed principal component analysis on the resulting growth vectors, which showed clear clustering of the strains by taxonomy and concentration of average loadings of sugars and acids along the first principal component.

### Genome analysis & Bioinformatics

Genomes for novel isolates were sequenced in multiple batches at the BioMicroCenter at MIT and at MiGS. We used SPAdes (35) with standard parameters to assemble draft genomes and checked genome completeness and contamination by CheckM (36). In cases, where first draft genomes were contaminated, we reisolated and sequenced until contamination was below 5%. In the final library, two strains remain with contamination >5%; those are excluded from all analyses.

Gene calling was performed by prodigal (37), genome annotation by eggnog v4.5 (mmseqs mode) (38) and CAZyme annotation by dbCANv2 (diamond mode) (39). Taxonomy assignment and creation of phylogenetic tree was performed by the standard workflow in GTDB-tk (40), followed by renaming of taxa falling into the NCBI clades Vibrionales and Alteromonadales (both assigned Enterobacterales by GTDB) to use the more familiar names for those clades.

Genome-scale metabolic models were created with CarveMe (41) using standard parameters. Growth on various carbon sources was simulated using cobrapy, using custom media emulating the mineral content of the experimental media (42).

Annotations and raw genome sequences were analyzed using Mathematica 12 (43) and R 4.2.0 (44). Trees were plotted using ggtree (45). Multiple-sequence alignment of amino-acid sequences for duplicated orthologs was performed using MEGA (46).

### KEGG pathway analysis

The KEGG ontology was used to define genes in different metabolic pathways. We analyzed the presence of KEGG pathways in three ways, for each strain and pathway:

- **Pathway completeness:** number of KOs with at least one copy present
- **Degree of duplication:** of those genes that are present, the total number of KOs that are present in more than one copy
- **Pathway coverage:** total number of KOs in the pathway, divided by the total genome size (number of KOs)

All correlations between pathway abundance measurements and SAP/GC content were corrected for multiple testing using the Bonferroni correction.

A complete list of KEGG orthologs and pathways considered for the analysis in the main text is given in SI Tables 4 and 5.

### Generalized linear model for SAP

To predict the SAP from genomic information, we developed two models. For the simple model described in the main text and used in Fig. 2c and d, we computed the total relative abundance of sugar and acid genes (unique genes in sugar and acid degradation pathways, see SI Table 5) from the eggnog annotations for each strain. We then used the two vectors of relative abundances 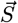 and 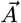 to predict the vector of 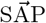 as

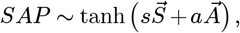

where the tanh function ensure that the predicted SAP values are between −1 and +1.

For the more complex model used in Fig. 3e, we used the same procedure but using the *pathway coverage* defined above for each of the 46 pathways given in SI Table 4.

### Progenomes

Reference genomes were downloaded from proGenomes and one genomes chosen randomly for each species (16). Genomes were annotated with eggnog; genomic GC content and the number of genes were extracted using custom R scripts. We computed pathway abundances for all genomes as described above and then predicted SAP by using those abundances as inputs into the linear model trained using our measured SAP.

### IJSEM trait database

Substrates used for growth for a large number of strains were obtained from a trait database assembled by Barberan et al. by text mining of articles in the International Journal of Systematic and Evolutionary Microbiology (15). The probability of growth for each substrate was measured by the number of entries within a particular range of genomic GC content that mentioned this substrate.

### Enrichment cultures

We analyzed relative abundance from enrichment cultures performed by Estrela et al. of 8 soil and leaf samples in 24 individual carbon sources (4 replicates each) (20). Briefly, 16S rRNA amplicons were sequenced and analyzed by dada2. We estimated genomic GC content for each ASV by matching taxon names with the SkewDB database contained genomic information on thousands of genomes. When exact species names did not match, we proceeded to the family level and finally order level, averaging over all representatives at that taxonomic level. Finally, we computed the abundance-weighted mean GC content per community accounting for the standard deviation of GC estimates.

## Data availability

Genomes and annotations are available under doi:10.17632/xfh8t8568g.1.

## ACKNOWLEDGEMENTS

We thank all members of the Cordero lab for discussions and feedback. This work was supported by the Simons Collaboration: Principles of Microbial Ecosystems (PriME) award number 542395. M.G. was supported by the Simons Foundation Postdoctoral Fellowship Award 599207.

## Supplementary figures

**Supplementary Figure S1.**
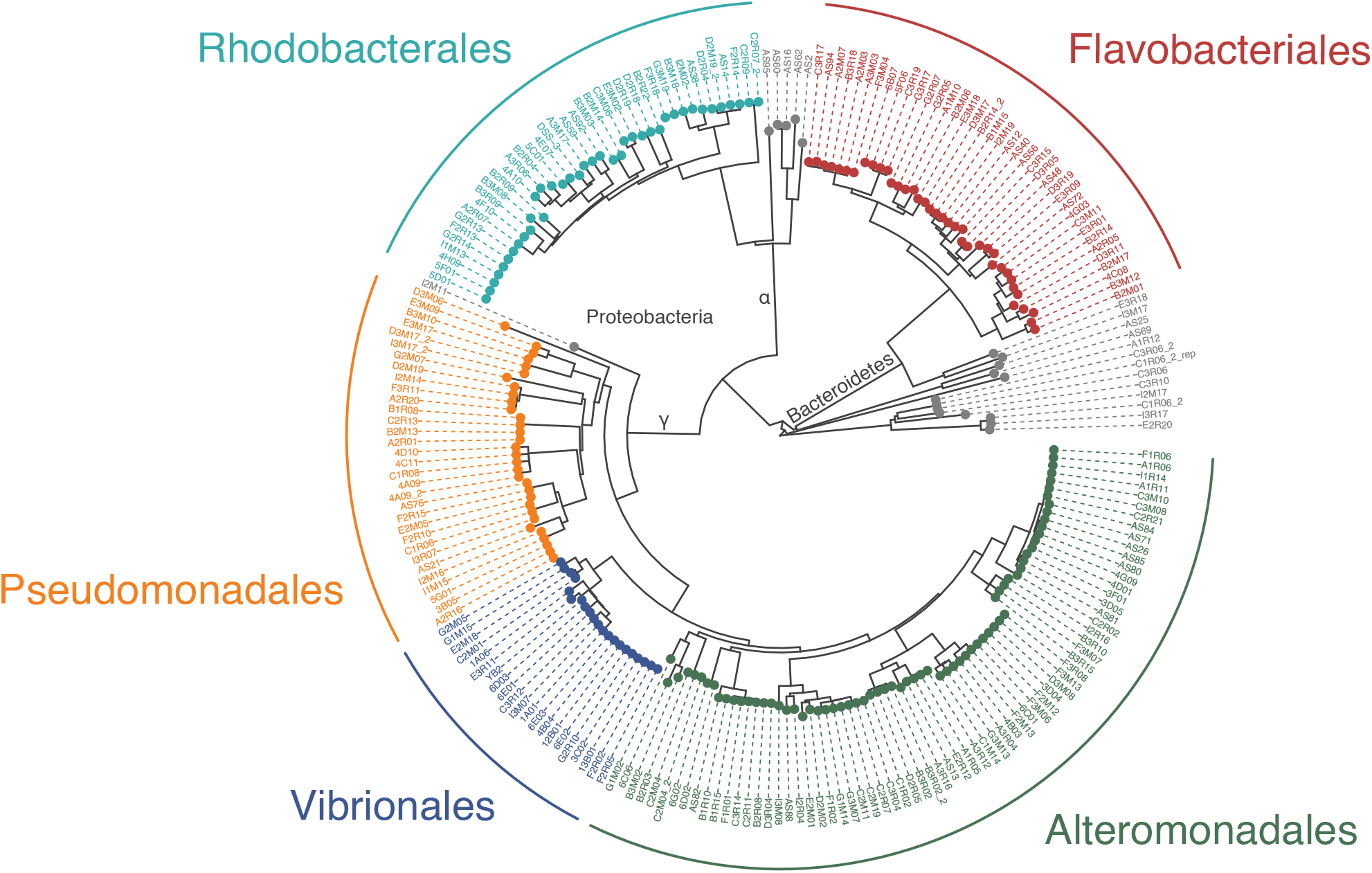
Phylogenetic tree, created using the GTDB-tk classify workflow using standard parameters, of all strains used in this study. A full list of all strains in given in SI Table 1.

**Supplementary Figure S2.**
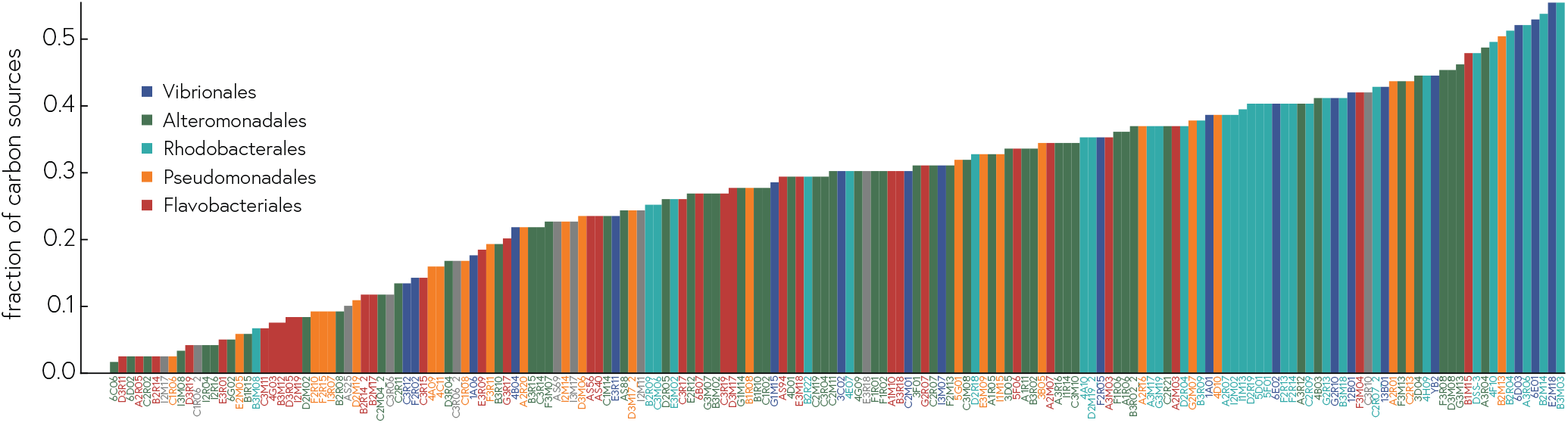
Number of carbon sources supporting growth per strain.

**Supplementary Figure S3.**
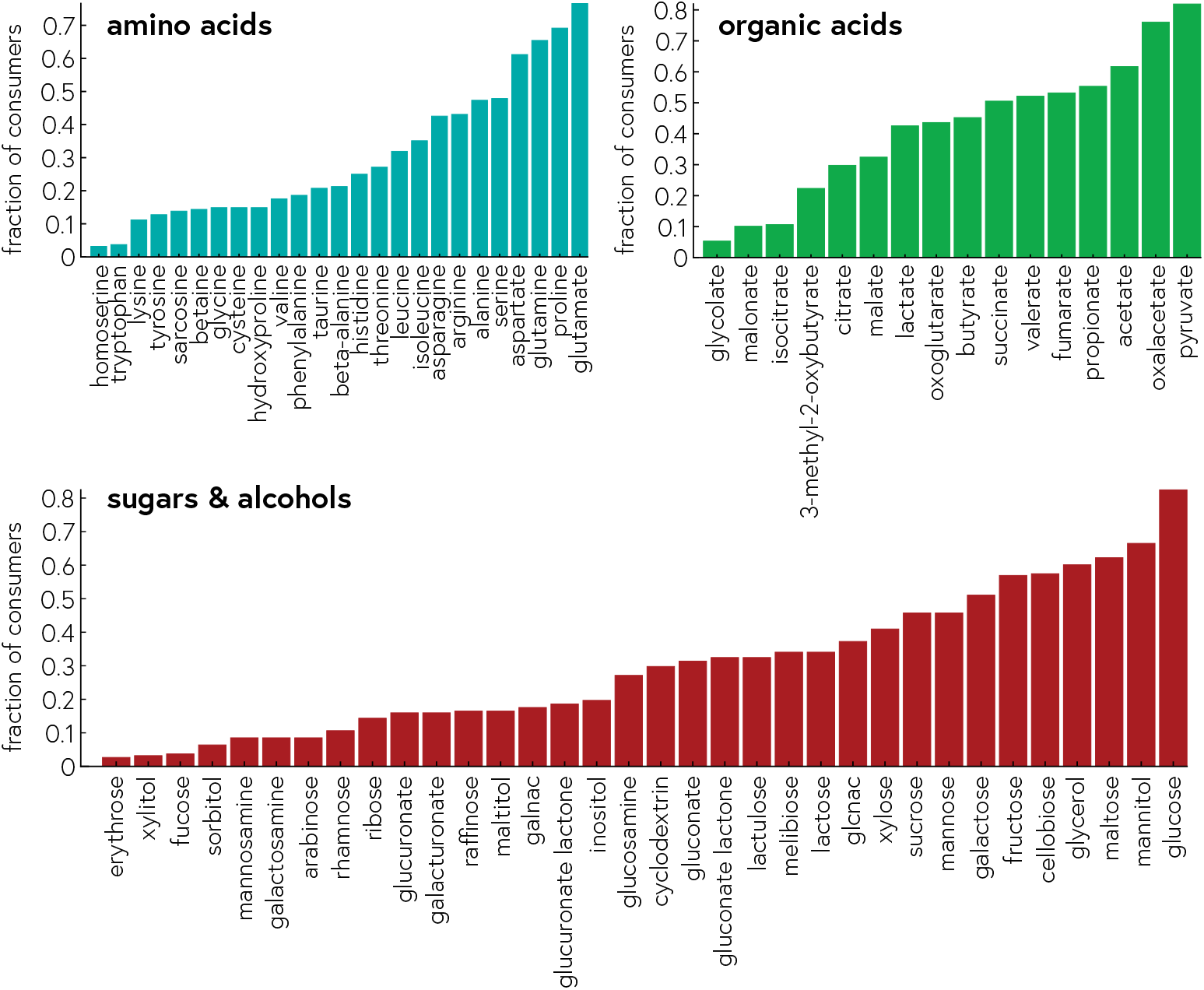
Fraction of all strains that were able to use a given substrate as sole carbon and energy source.

**Supplementary Figure S4.**
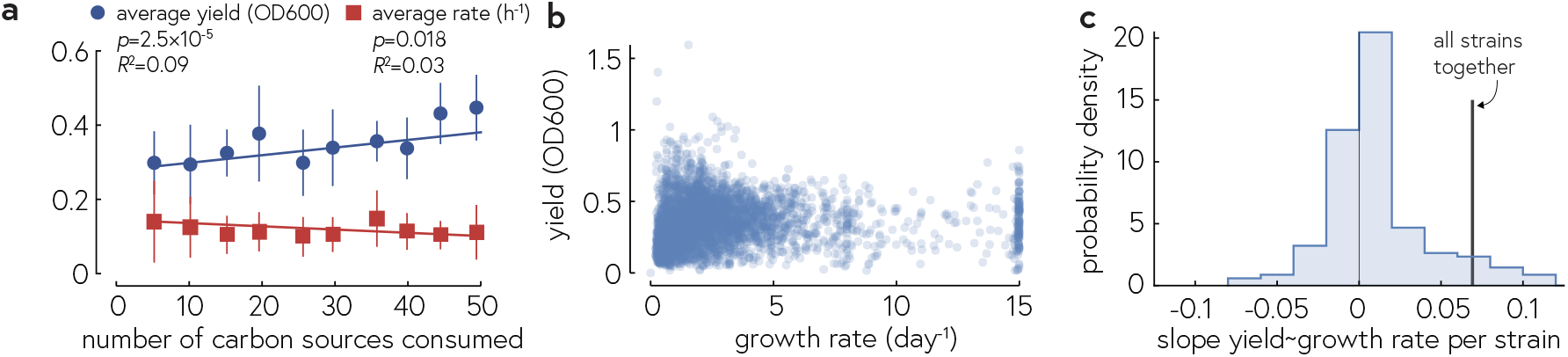
Lack of strong correlation between number of carbon sources that support growth, growth rate, and yield. a) Average yield (blue dots) and rate (red squares) binned by the number of carbon sources that supported growth. More generalist species (more carbon sources consumed) achieve slightly higher average yield, but the effect size is likely not practically relevant. b) For each condition (substrates*×* strain, we plot the growth rate and yield, which are very slightly positively correlated (p = 2 *×* 10^*−*6^, R^2^ = 0.005). c) Linear slopes for the per-strain regression of yield with growth rate; only 3/186 strains exhibited a statistically significant. The vertical line corresponds to the slope of the regression in panel b.

**Supplementary Figure S5.**
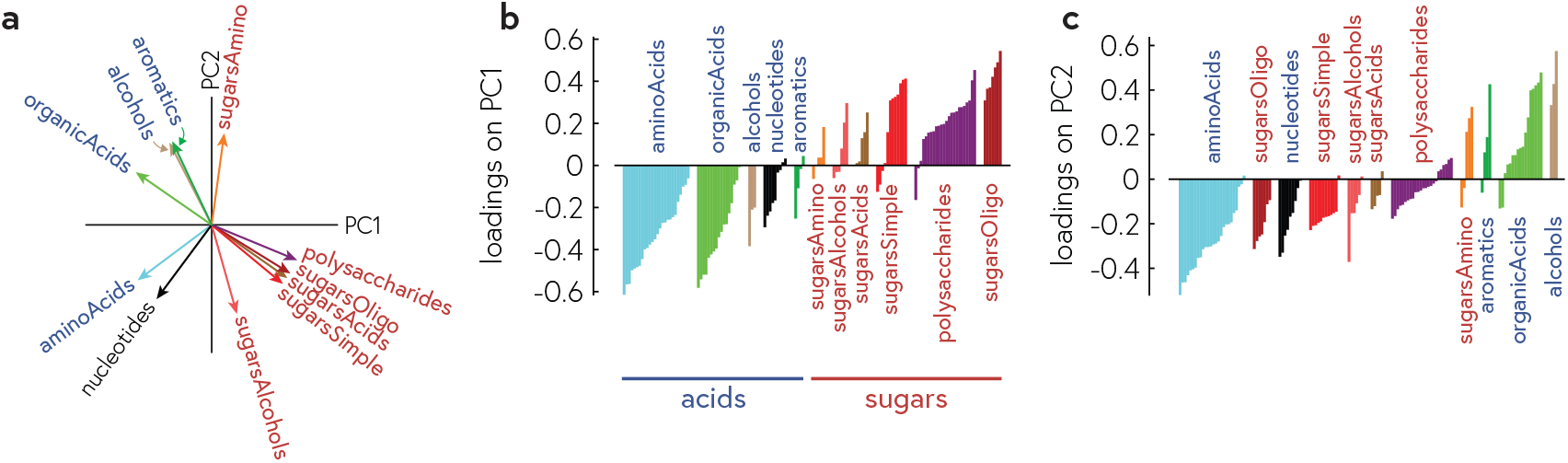
Detailed loadings of all substrates in the Principal Component Analysis in Fig. 1. a) Averaged loadings of fine-grained categories of substrates, normalized to unit length. b) Individual loadings per substrate for each principal component (PC). Note how all acids have negative loadings on PC1 but all but one organic acids switch sign on PC2 relative to amino acids.

**Supplementary Figure S6.**
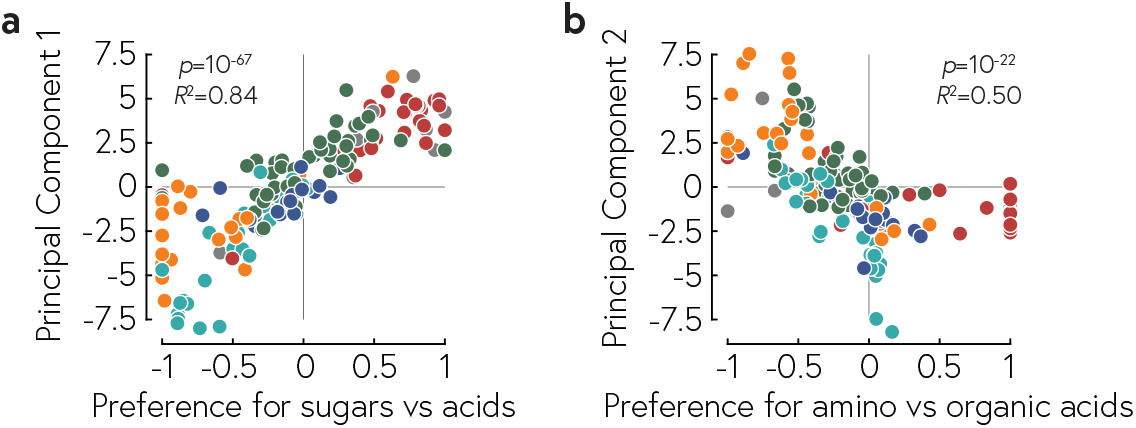
Scatter plots of the first principle component vs the Sugar-Acid preference (SAP) as defined in the main text, and the second principle component vs the Amino Acid-Organic Acid preference (OAP) defined analogously.

**Supplementary Figure S7.**
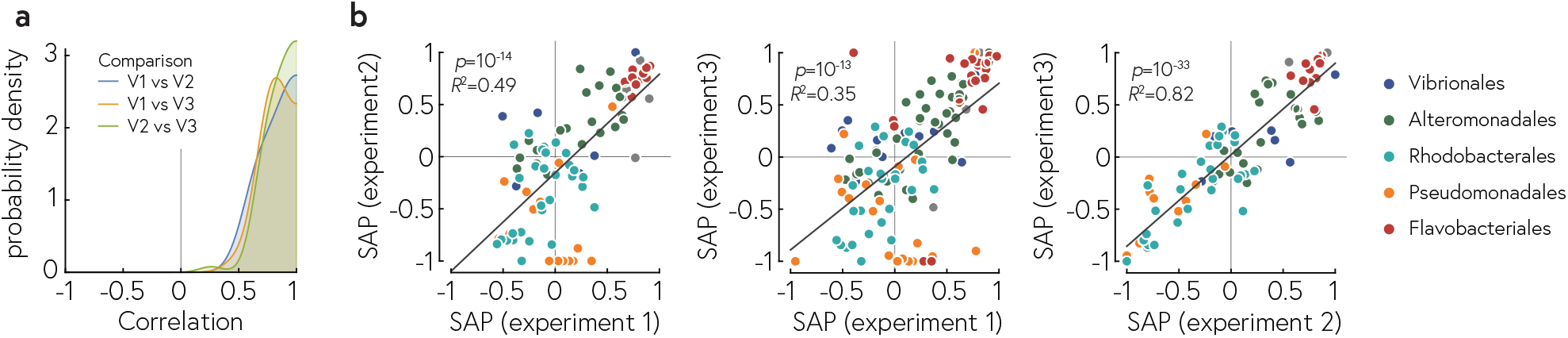
Reproducibility between experiments. a) Correlation coefficients between all three experiments (V1, V2, V3; V3 is the experiment primarily discussed in the main text), correlating the ability to grow on a given substrate across all strains. b) Scatter plots of the SAP measured for each strain between all three replicate experiments.

**Supplementary Figure S8.**
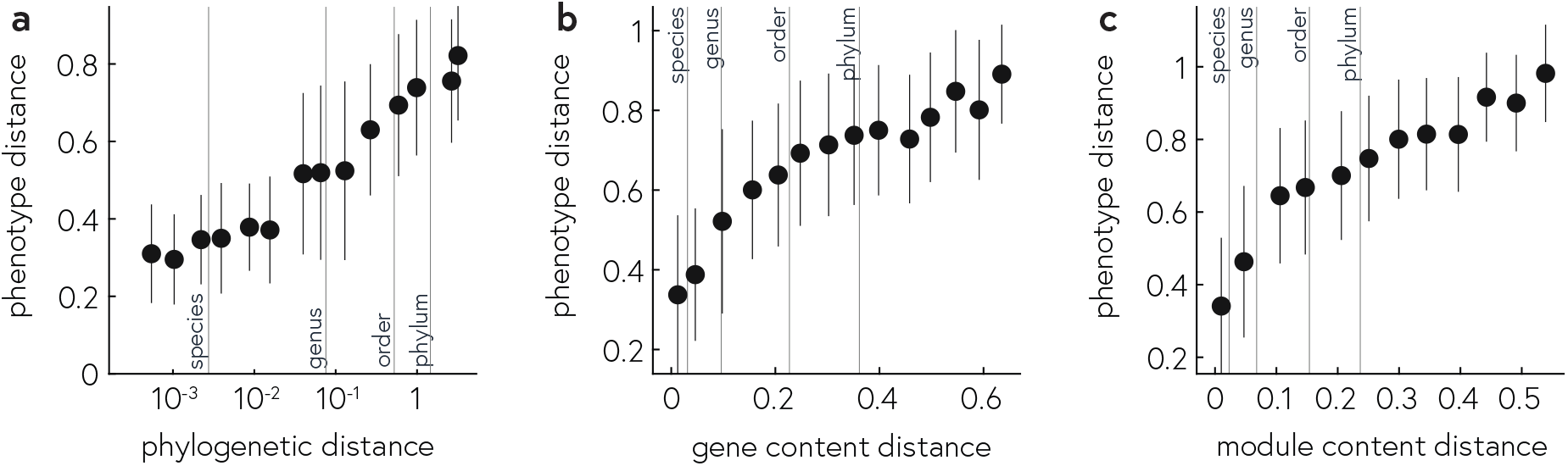
Phenotype distance, defined as the cosine distance between consumption vectors, as a function of genomic distance between pairs of strains, where the genomic distance is the GTDB-tk distance (a) or the Bray-Curtis distance between gene content (panel b, based on copy numbers of KEGG KO) or module content (panel c, based on abundance of KEGG modules).

**Supplementary Figure S9.**
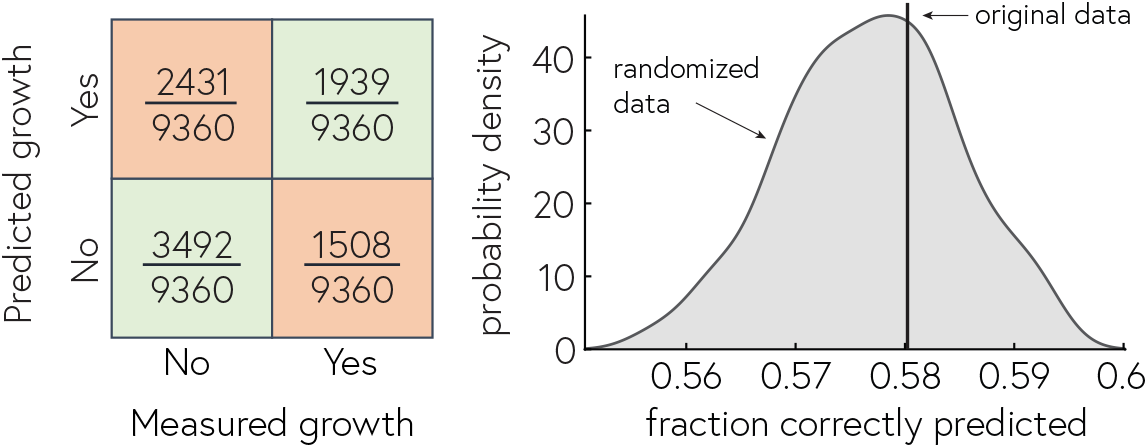
Comparison between measured and predicted growth on individual substrates. Predicted growth was derived from FBA simulations of genome-scale metabolic models created using CarveMe using standard parameters (no gapfilling). This procedure yielded 58% correct predictions (vertical line), which was within the range of correct predictions achieved when the comparison was performed with shuffled labels (distribution, obtained by shuffling labels 1000 times, each time measuring the proportion of correct predictions).

**Supplementary Figure S10.**
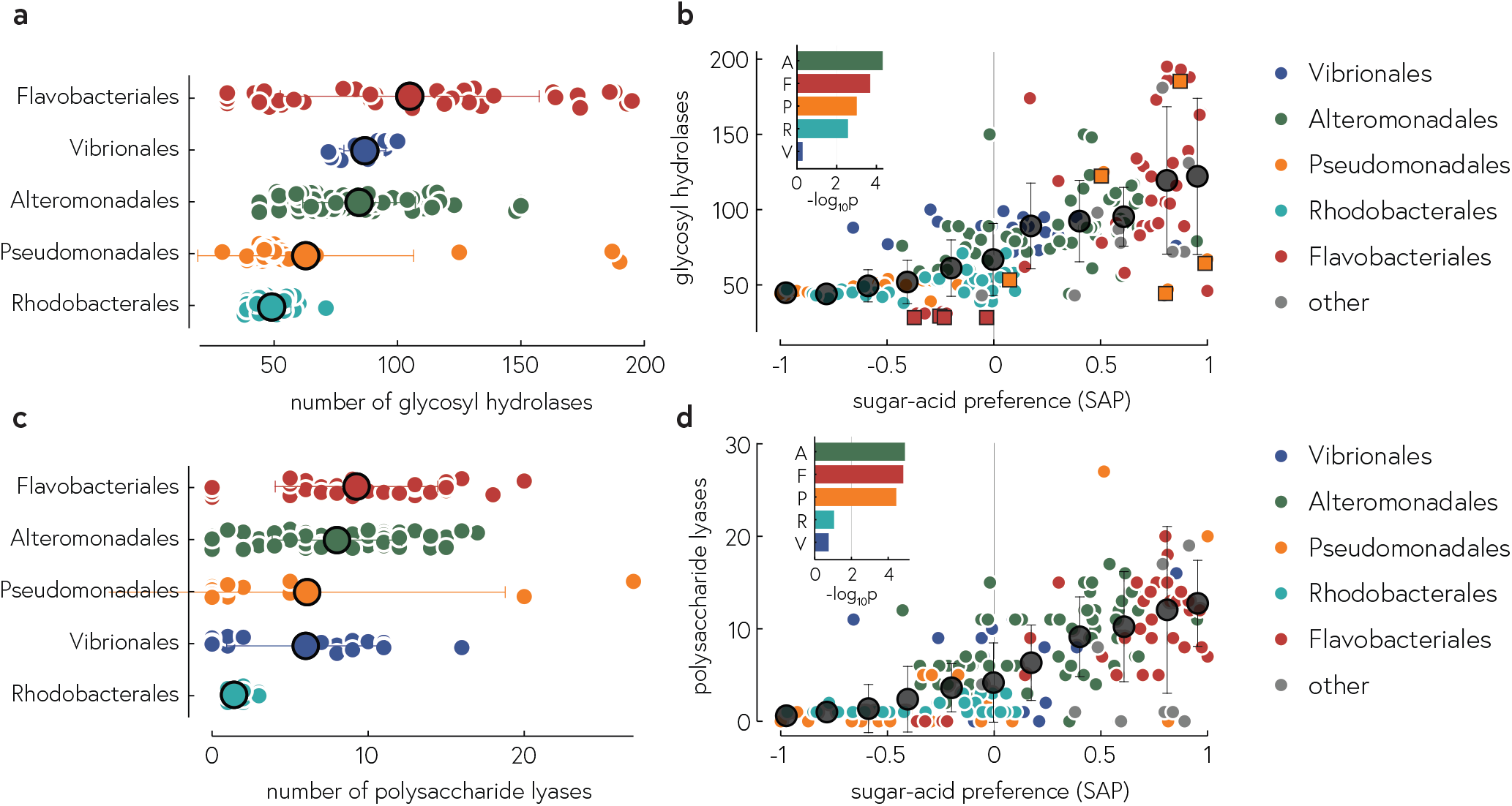
Number of CAZymes (glycosyl hydrolases, top; polysaccharide lyases, bottom) and their correlation with sugar-acid preferences (right panels). The insets show *−* log_10_ p per order, the negative log_10_ of the p-value obtained from linear regressions of CAZyme number with SAP within each order. *−* log_10_ p > 2 (vertical line) corresponds to a significant correlation at the 5% level, Bonferroni correcting for multiple testing. The square symbols in panel b correspond to the squares in Fig. 1d, i.e., the flavobacteriales and pseudomonadales strains with atypical phenotypes for their taxonomy, which have fewer/more CAZymes than their close relatively, respectively (see main text).

**Supplementary Figure S11.**
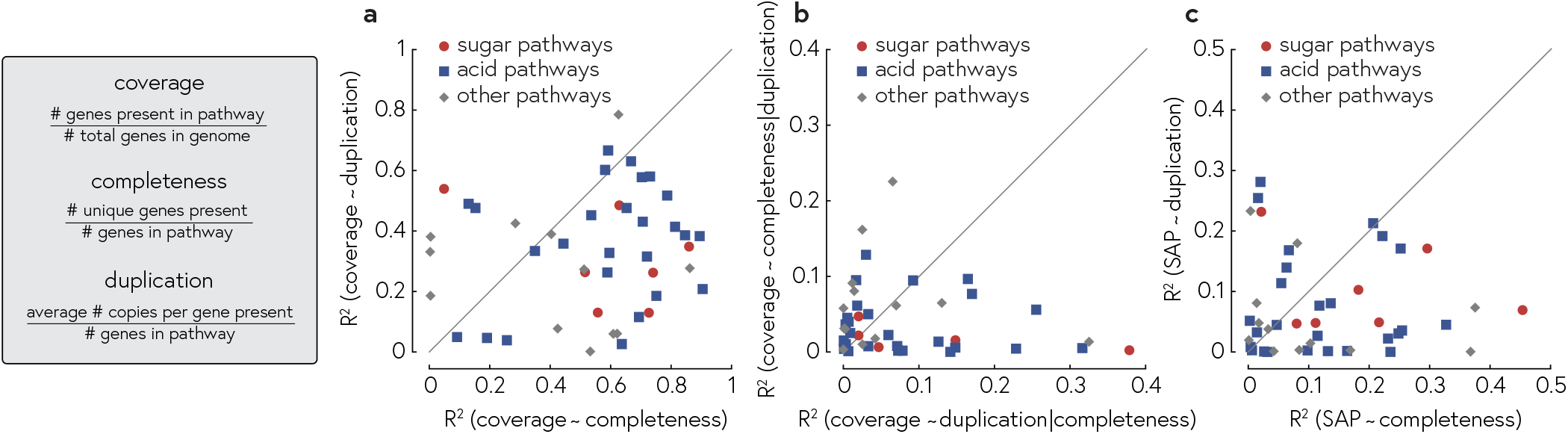
Three measures of pathway abundance, as defined in the Methods, and their interrelations. a) Predicting coverage from completeness (linear model) generally yields higher quality fits than predicting coverage from duplication. b) After correcting for completeness, duplication tends to explain more of the residuals than completeness does after correcting for duplication. Neither duplication nor coverage of any individual pathway correlated very strongly with SAP, and whether duplication or coverage of a given pathway was more predictive of SAP depended on the pathway.

**Supplementary Figure S12.**
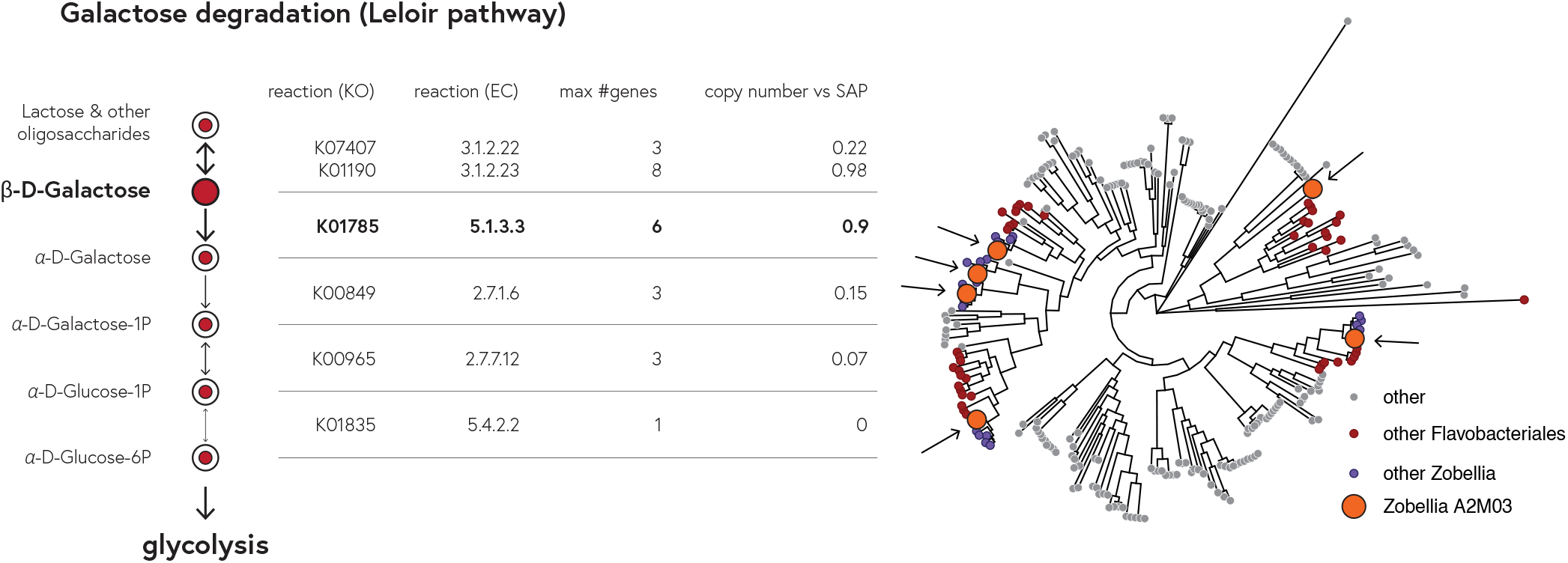
Illustrating the concept of functional duplication on the example of the galactose degradation pathway (KEGG pathway ko00052). Shown is the central part of the pathway that converts lactose and other oligosaccharides first to β-D-galactose, which is transformed through multiple steps to α-D-glucose-6-phosphate, which then enters glycolysis. For some reaction, we found multiple orthologs in the same strains (e.g., up to 6 orthologs of K01785 (galM, aldose 1-epimerase, EC:5.1.3.3). These ortholog are not pure duplication, as illustrated by the tree on the right. The tree is based on a multiple-sequence alignment of all sequences annotated K01785 across all strains. We have highlighted the 6 copies found in the *Zobellia* strains A2M03, which are spread around the tree and often grouped with orthologs found in distantly related species. In fact, across all highly duplicated orthologs (maximum number of orthologs per strains at least 6), the pairwise distance (computed from the tree using the *cophenetic*.*phylo* function in the *ape* library in R), was about equally likely to be greater between orthologs in the same strain relative to orthologs in different strains, as it was to be smaller. Thus, “duplicated” orthologs in a strain likely represent functional variants of different evolutionary origin.

**Supplementary Figure S13.**
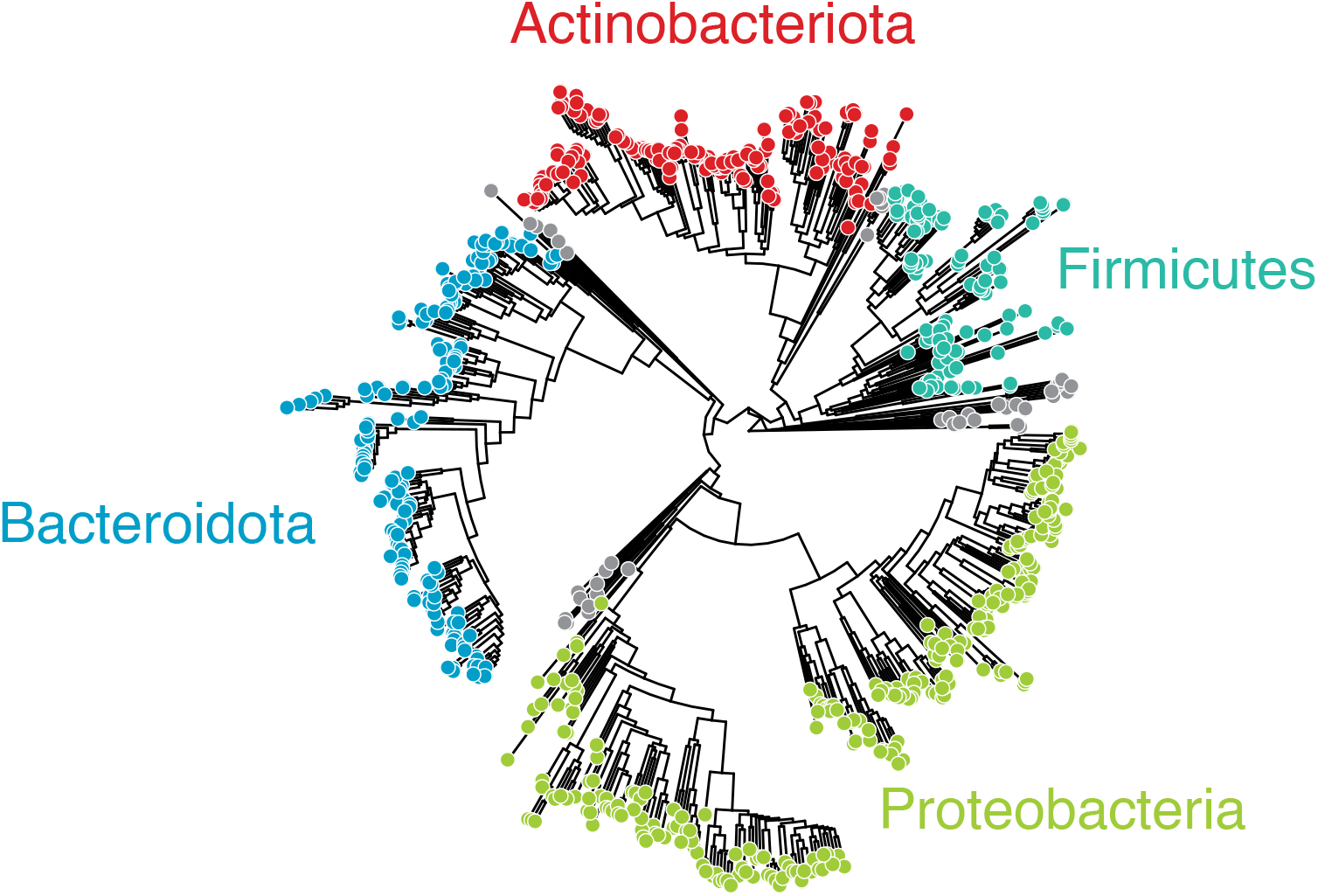
Phylogenetic tree based on GTDB-tk of species contained in the IJSEM trait database (15) as well as progenomes (by species name) (16). Note that two phyla, Actinobacteriota and Firmicutes, are not at all represented in our strain library.

**Supplementary Figure S14.**
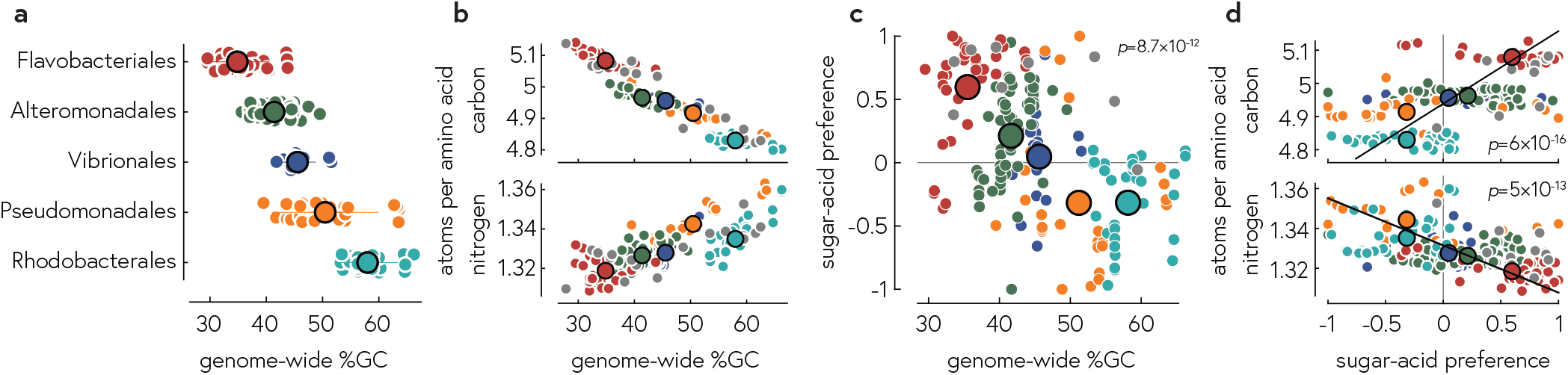
Genomic GC content and consequences for nutrient requirements. a) GC content is relatively conserved at the order level across our strain library. b) GC content predicts the carbon and nitrogen requirements per coded amino acid. c) Same data as Fig. 3a without binning: GC content is correlated with genomic GC content across the whole set of strains, but not within orders. Because of the correlation of GC content with both nutrient requirements and SAP, SAP is positively/negative correlated with the number of carbon/nitrogen atoms per coded amino acid.

**Supplementary Figure S15.**
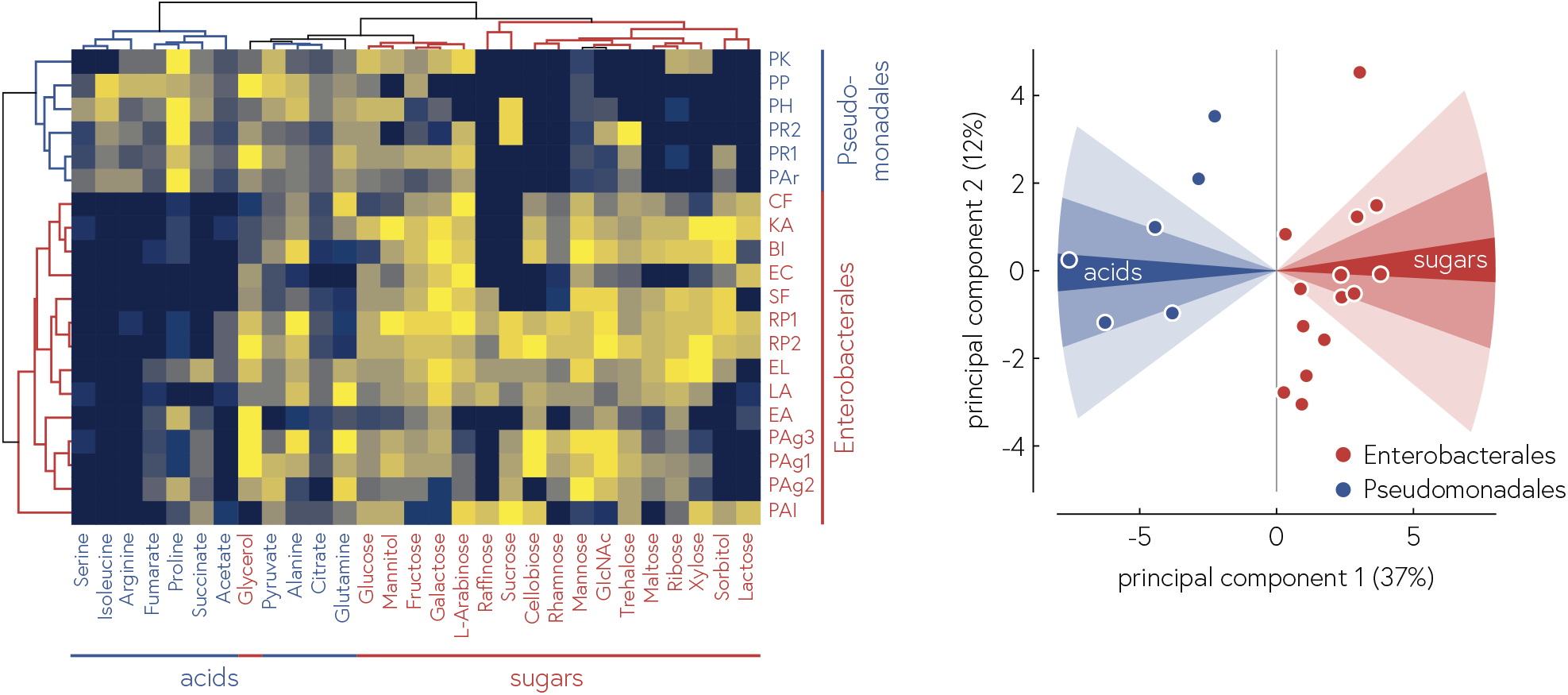
Re-analysis of data from Kehe et al. (34). The heatmap corresponds to SI Fig. S2 in Ref. S15, but with rows and column sorted by cosine similarity. Principal component analysis of this matrix shows the clustering of the two taxonomic orders and their alignment with the average loadings of acids and sugars.

**Supplementary Figure S16.**
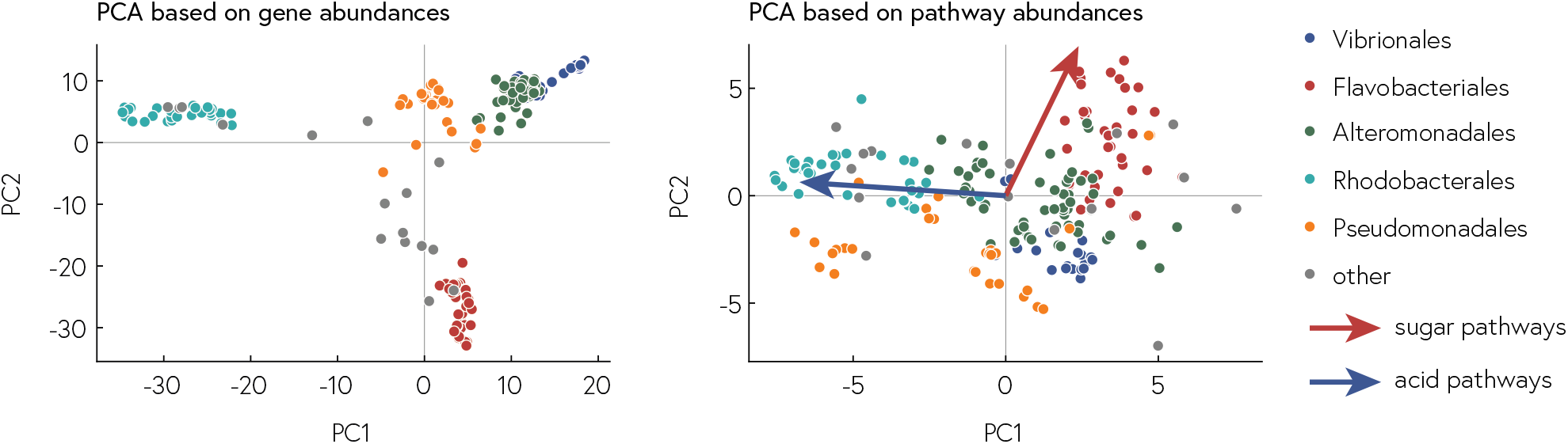
Principal component analysis (PCA) of genome content reproduced taxonomic structure. a) PCA based on relative abundances of individual genes (see SI Table 3 for a list of the 2505 genes in 46 pathways used for the analysis) in central metabolism showed clear clusters for the different orders. b) PCA based on relative abundances of the same 46 pathways, with the average loadings of sugar and acid pathways.

**Supplementary Note 1: Biological replicate experiments**

Following pilot experiments, we performed variants of the experiments three times, with slight alterations to strains, substrates, and procedure. The final experiment 3 is described in the main methods. Substrates that did not support growth in any species in experiment 3 are indicated in SI Table 1. Additional substrates tested in experiments 1 and 2 are listed below.

**Experiment 1**. Experiment 1 was performed before draft sequences were obtained for all strains and therefore contained contaminated strains which were subsequently removed from the analysis and replaced with either reisolated or new strains; the final set of strains is given in SI Table 2.

The following substrates were used in experiment 1 but were not included in the final experiment because they yielded no or very little growth: sodium formate, L-lyxose, ethylene glycol, maleic acid, L-sorbose, acetaldehyde, norvaline, and PABA (para-aminobenzoic acid).

The inoculation procedure for experiment 1 was as follows: after 4 days of growth in MB at room temperature, strains were transferred into fresh media in 384 well plates using a metal pinning tool (VP 384, V&P Scientific) that was cleaned between plates by first washing in water, then pure ethanol. The pinning tool was then flame sterilized and cooled for 1 minutes.

**Experiment 2**. Experiment 2 was performed similarly to experiment 1, except that disposable pinning tools (VP 248, V&P Scientific) were used for each plate and immediately discarded.

The following substrates were used in experiment 2 but were not included in the final experiment because they yielded no or very little growth: sodium glyoxylate, L-lyxose, ethylene glycol, norvaline, and PEP.

